# Identification and characterization of two CRISPR-Cas systems associated with the mosquito microbiome

**DOI:** 10.1101/2023.03.22.533747

**Authors:** Shivanand Hegde, Hallie E. Rauch, Grant L. Hughes, Nikki Shariat

## Abstract

The microbiome profoundly influences many traits in medically relevant vectors such as mosquitoes, and a greater functional understanding of host-microbe interactions may be exploited for novel microbial-based approaches to control mosquito-borne disease. Here, we characterized two CRISPR-Cas systems in a novel bacterium, *Serratia* Sp. Ag1, that was isolated from the gut of an *Anopheles gambiae* mosquito. Two distinct CRISPR-Cas systems were identified in *Serratia* Ag1, CRISPR1 and CRISPR2. Based on *cas* gene composition, CRISPR1 is classified as a Type I-E CRISPR-Cas system and has a single array, CRISPR1. CRISPR2 is a Type I-F system with two arrays, CRISPR2.1 and CRISPR2.2. RT-PCR analyses show that all *cas* genes from both systems are expressed during logarithmic growth in culture media. The direct repeat sequence of CRISPRs 2.1 and 2.2 are identical and found in the arrays of other *Serratia* spp, including *S. marcescens* and *S. fonticola*, whereas CRISPR1 was not. We searched for potential spacer targets and revealed an interesting difference between the two systems: only 9% of CRISPR1 (Type I-E) targets are in phage sequences and 91% are in plasmid sequences. Conversely, ~66% of CRISPR2 (Type I-F), targets are found within phage genomes. Our results highlight the presence of CRISPR loci in gut-associated bacteria of mosquitoes and indicate interplay between symbionts and invasive mobile genetic elements over evolutionary time.

**Data Summary:** There is no supporting external data generated for this work.

## Introduction

Host associated microbes play a crucial role in the physiology, diseases, and immunity of their host. In mosquitoes, gut associated microbes profoundly affect their host and these altered phenotypes influence vectoral capacity and vector competence (1–6). Bacteria are abundant constituents of the gut microbiome of mosquitoes (7–10), however metagenomic studies have also found bacteriophage associated with these vectors (8–12), and it would be reasonable to expect interplay between these microbes given their co-ocurance. While microbe-microbe interactions within the gut alter bacterial community structure and colonization (13, 14), less is known regarding the interactions between bacterial communities and bacteriophages, however signatures of these encounters can be inferred from bacterial genomes.

Clustered regularly interspaced short palindromic repeats (CRISPR)-Cas systems are present in approximately 45% of sequenced bacterial genomes, and 90% of archaeal genomes (15). In their canonical function, they act as a small RNA-driven adaptive immune system that provides defense against exogenous nucleic acids, namely bacteriophage and plasmids (16, 17). CRISPR-Cas systems have two components, a suite of *cas* genes and a CRISPR array (18). The latter comprise direct repeat sequences ranging from 21-48 nucleotides in length that separate highly variable spacer sequences of similar lengths (19). Spacers are commonly derived from foreign nucleic acids and are added in a polar manner to the CRISPR array, with the newest spacers being found closest to the leader sequence. CRISPR immunity takes place in three distinct steps: first new spacers are acquired and added to the array as the prokaryote adapts to a new invader (16, 20, 21). Second, the array is transcribed, and the resulting transcript processed to produce mature CRISPR RNAs (crRNAs) (21, 22). Third, the crRNA guides an endonuclease to its complementary target nucleic acid, thereby resulting in degradation, or interference, of the target (17, 21). Various *cas* gene products are required for each of these steps. CRISPR-cas systems can be separated into two distinct classes and into further subtypes, depending on the complement and organization of *cas* genes (23). Class 1, Type I systems are defined by the inclusion of Cas3 as the effector endonuclease responsible for cleaving target DNAs. Within Type I, there are seven subtypes, I-A - I-G (24–26). Subtypes I-E (e.g., found in *Escherichia coli* and *Salmonella enterica)* and I-F (e.g., found in *Yersinia pseudotuberculosis* and *Pectobacterium atrosepticum)* differ slightly from each other. Type I-E has a distinct Cas2 protein, whereas in Type I-F, Cas2 and Cas3 form a chimeric protein. Further, Type I-F systems also lack Cas11, which forms part of the Type I-E effector complex (22, 27). In the Enterobacteriaceae family, CRISPR-cas systems almost exclusively belong to either Type I-E or Type I-F (22, 27).

In addition to their well-characterized role in prokaryote adaptive immunity, alternative functions have also been attributed to some CRISPR-Cas systems (20). These include roles in biofilm formation, host avoidance, and symbiosis and highlight the important biological roles of these systems in pathogenic bacteria, as well as other bacterial species (20, 28). Given this, and the recent explosion in genome editing capabilities of Cas genes, there is a drive to discover new CRISPR-Cas systems in a wide array of prokaryote genomes. CRISPR-cas systems in host associated microbiomes has mainly been examined in the context of human and plant microbiomes (29–32) while invesigations in invertebrates are lacking. Studies focused on bacteria that play integral roles in the human microbiome have revealed important roles for CRISPR-Cas in viral resistance and mitigation of foreign genetic material (32–35). Though CRISPR-cas technology has been applied for genome editing of mosquito vector host and their microbiome (36–38), characterizing native CRISPR loci in gut bacteria of mosquitoes has not been attempted so far.

To determine interactions between gut associated bacteria of mosquitoes and bacteriophage over evolutionary time, we examined the genomic signature of CRISPR-Cas systems in Ag1, a *Serratia* strain previously isolated from *Anopheles gambiae* mosquitoes (39). We found Ag1 harbors two Type I CRISPR systems and further classification revealed they belong to subtype I-E and I-F. We also examine the origins of the spacer region, thereby identifying past infections of the bacterial host, and characterized the expression of the Cas genes. Our results indicate the presence of CRISPR-Cas systems in symbiotic bacteria associated within invertebrates and highlight the complexity of microbial interactions within the mosquito gut.

## Materials and Methods

### Culturing and nucleic acid isolation

Origin of the bacterial isolates *Serratia sp.* Ag1 and *Serratia sp.* Ag2 (JQEI00000000 (Serratia sp. Ag1) and JQEJ00000000 (Serratia sp. Ag2)) used in this study were described before (39). Total genomic DNA was isolated from overnight cultures of Ag1 and Ag2 using the Genome Wizard kit (Promega, WI), following the manufacturer’s protocol. DNA pellets were resuspended in 200 μl of molecular grade water and stored at −20°C. Bacterial strains were cultured in LB broth to log phase and to stationary phase and total RNA was isolated using TRIzol (Life Technologies, CA) and resuspended in 20 μl molecular grade water. RNA was treated with 1-unit Dnase (Life Technologies, CA) and re-isolated with TRIzol. Pellets were resuspended in 20 μl molecular grade water and stored at −20°C.

### RT-PCR expression analyses

A total of 100 ng total RNA was used to generate cDNA in a 20 μl reaction using a qScript mastermix (QuantaBio, MA) that contained random hexamers. Reverse transcription was performed in a PCR machine with the following parameters: 22°C for 5 minutes, 42°C for 30 minutes, 85°C for 5 minutes, and 4°C hold. For a non-RT control, reactions were set up in duplicate but without RT enzyme. The cDNAs were diluted 1:10 and 2 μl of each used for subsequent PCR reactions with one unit of Taq polymerase (New England Biolabs, MA), 200uM dNTPs (New England Biolabs, MA) and 1x Standard Taq Polymerase buffer in a 25 μl reaction. The primers used for RT-PCR analysis of *cas* genes are listed in Table 1. Following initial denaturation for three minutes at 95°C, PCR conditions were as follows: 20 cycles (16S control PCR) or 25 cycles (cas genes) of 95°C for 30 seconds, annealing at 57°C for 30 seconds, and extension at 72°C for 30 seconds. A total of 5 μl of the PCR reaction was imaged by gel electrophoresis.

**Table 1:**
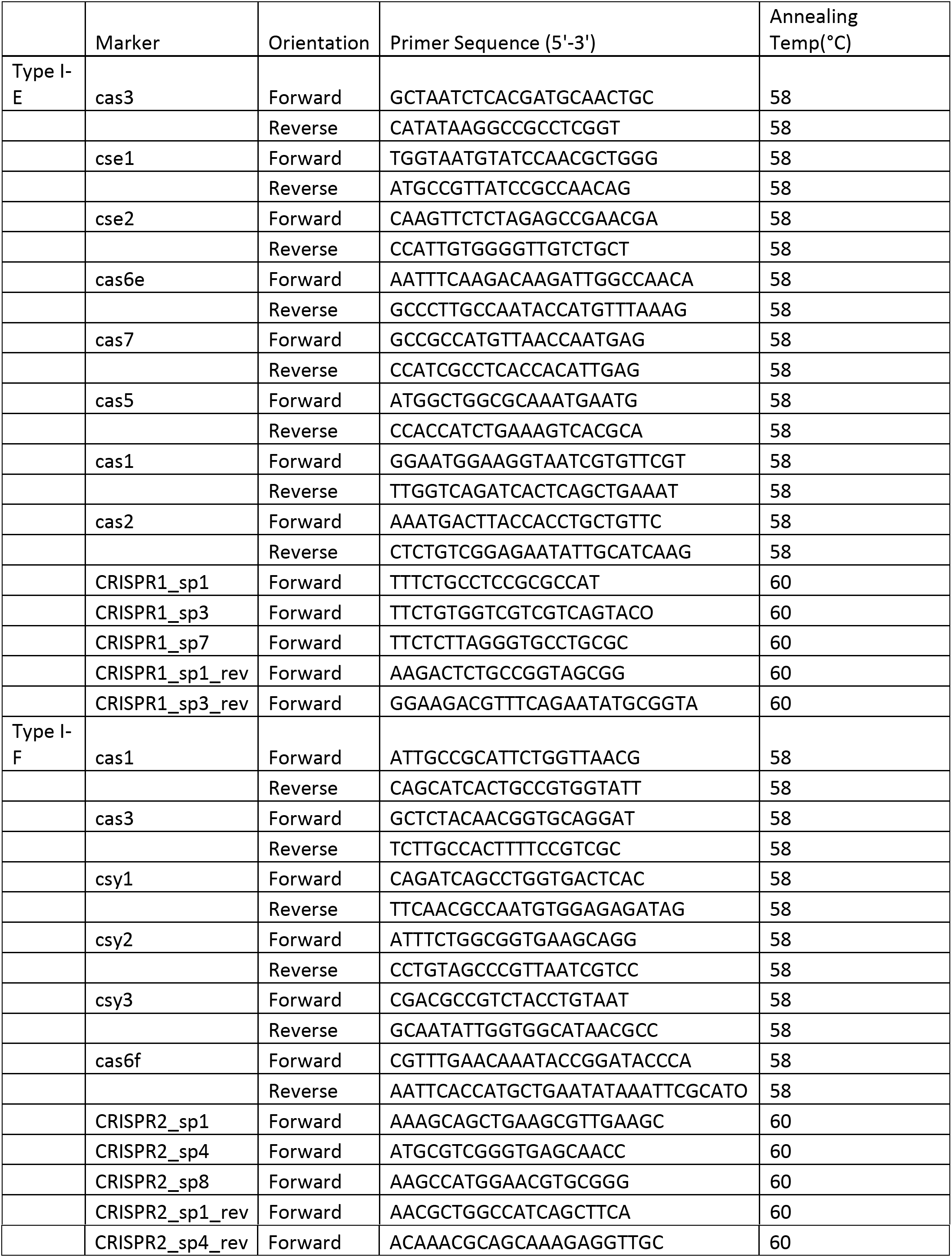
Primers used in this study

### Identification of CRISPR loci, phylogenetic analyses, and spacer identification

The assembled Ag1 genome was analyzed using CRISPR-Finder (40) to identify both the CRISPR arrays and the *cas* genes. Spacers were extracted from the arrays and analyzed using an Excel-based macro (41). CRISPR Target (42) was used to identify putative spacer matches. We considered matches to be 24/32 or 24/33 nucleotides for the Type I-E and I-F spacers, respectively. For phylogenetic analyses, the coding sequence of both *cas3* genes were translated and BLAST was used to find the top 20 similar sequences from different species. These amino acid sequences were used in MEGA7 to build phylogenetic trees with a bootstrap value of 1000 (43).

## Results

We identified two Type I CRISPR-Cas systems in Ag1 and termed them CRISPR1 and CRISPR2. The former has a single CRISPR array and is of the Type I-E subtype of CRISPR-Cas systems (Figure 1a), with direct repeats and spacers that are 28 and 33 nucleotides long, respectively. The CRISPR2 has a *cas* operon associated with the Type I-F subtype, and there were two CRISPR arrays associated with this system, which we termed CRISPR2.1 and CRISPR2.2. The direct repeats and spacers in both arrays are 28 and 32 nucleotides in length, respectively. The Type I-E repeat sequences fall under Cluster 2 and the Type I-F direct repeat sequences fall under Cluster 4 (18, 25, 44). Spacer composition of the three CRISPR arrays in Ag1 was analyzed and the spacer content of each array was distinct (Figure 1b). CRISPR2.1 was the longest array and contained 26 different spacers.

**Figure 1.**
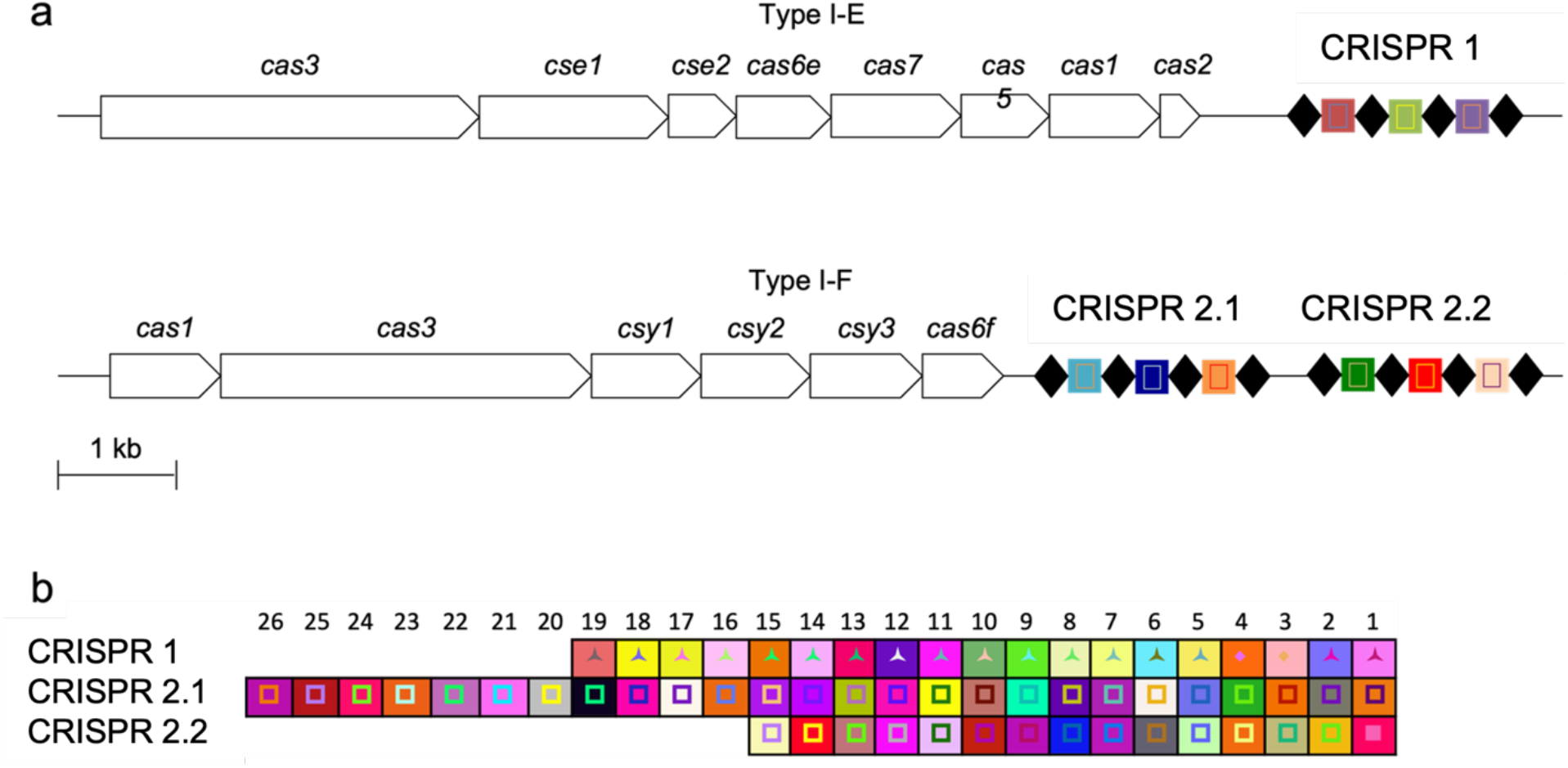
Organization and expression of the Type I-E and Type I-F CRISPR-cas systems of *Serratia* sp. Ag1. **a.** All *cas* genes are shown in the forward orientation. Direct repeats in the CRISPR array are shown as black diamonds while the spacer sequences are represented by colored squares. The *cas* genes are scaled to the 1 kb bar shown in the bottom left. **b.** Spacer composition of the three CRISPR arrays in Ag1. The unique combination of the background color and the shape and color in the foreground represents a single spacer sequence. The three-point star represents a spacer that is 33 nt in length. The inner square represents a 32 nt spacer. The oldest spacer, #1, is shown to the far right, while the most recently acquired spacer is shown on the far left. The invariant direct repeats have been removed for clarity.

Using the cas3 protein sequence from each CRISPR-Cas system, we identified similar protein sequences from other bacterial species and examined their phylogeny. We found a single match to another *Serratia* sp. Ag2, which is closely related to Ag1 (45) (figure 2). Otherwise, we did not find any other *Serratia* spp. whose cas3 matched closely to the cas3 of CRISPR1, suggesting that type I-E system is not broadly present in other *Serratia* spp. The Type I-E cas3 was closely related to *Dickeya* spp. and *Klebsiella* spp. and overall, there was little divergence among the Type I-E cas3 proteins compared to those from the Type I-F subtype (Figure 2). Conversely, we found several *Serratia* spp. that contained cas3 protein sequences of the Type I-F subtype, however the sequence from Ag1 was more closely related to some *Yersina* spp. than those *Serratia* spp.

**Figure. 2:**
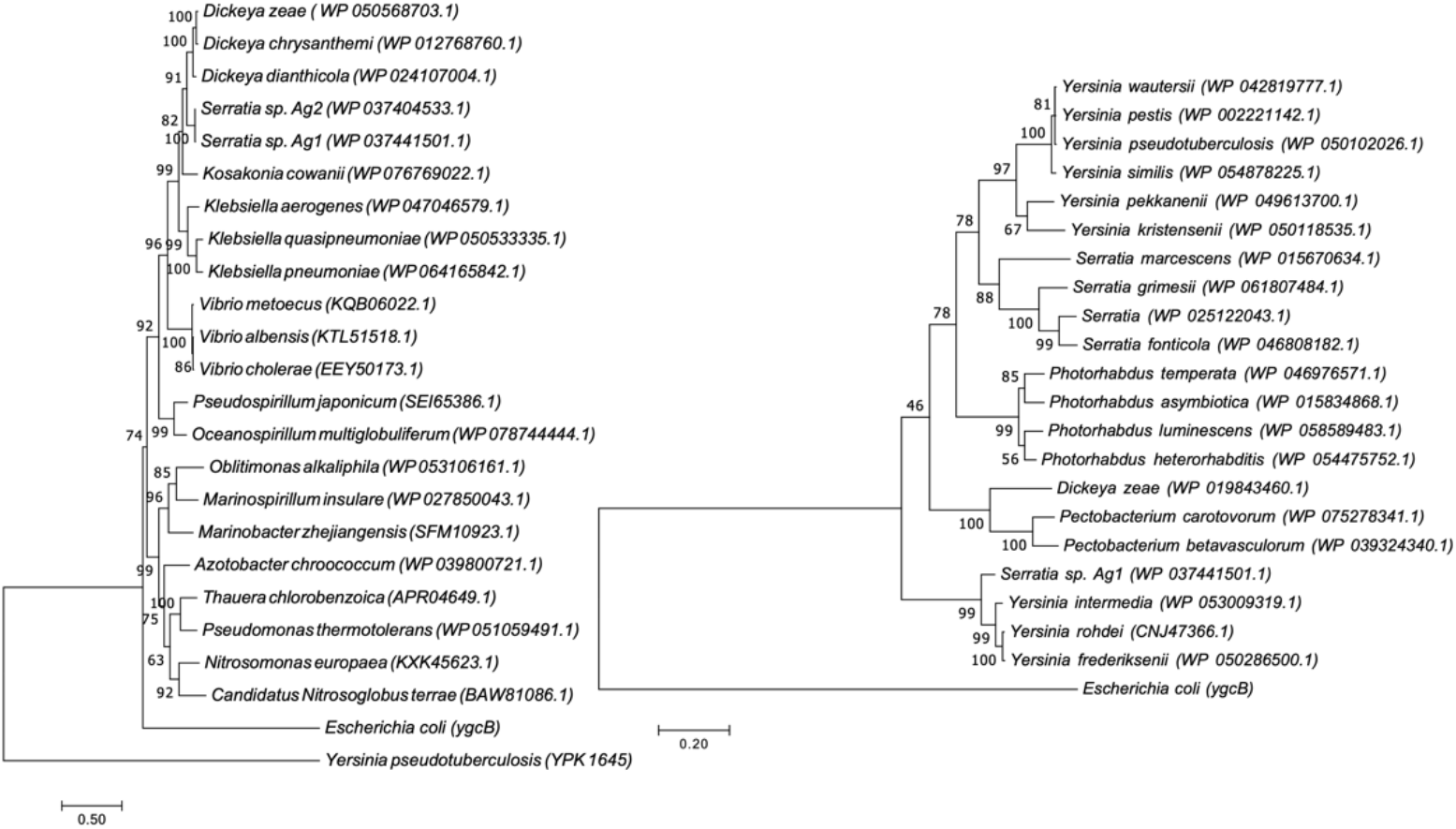
Phylogenetic analyses of Type I-E and Type I-F Cas3 from *Serratia* sp. Ag1. Phylogenetic trees show analyses of Cas3 with the top 20 closes BLAST-p hits for both trees. Maximum likelihood trees based on the relevant Cas3 protein are shown with a bootstrap value of 1000. *E. coli* is included as a representative of the Type I-E subtype, and *Yersinia pseudotuberculosis* is included as a representative of Type I-F.

We analyzed the CRISPR spacers to determine whether they matched to any exogenous nucleic acids and found a greater number of matches to plasmid and bacteriophage (including prophage sequences) sequences in CRISPR2.1 (50%, 13/26 spacers had matches) and CRISPR2.2 (53%, 8/15) than in CRISPR1 (32%, 6/19) (Figure 3, Table 2). In both CRISPR2.1 and 2.2, phage targets accounted for the most hits, constituting two thirds of the identified targets (Figure 3). Conversely, although CRISPR1 had fewer spacer targets that we could identify, most targets were of plasmid origin (Figure 3). When we increased the stringency of the matches to 85% (28/33 nucleotides for the CRISPR1 array, 27/32 for CRISPR2 arrays), we reduced the number of hits significantly. Of the remaining 19 spacer targets, only one matched to a spacer in CRISPR1 and 16 spacers matched to a phage target. Expression of the *cas* genes from both subtypes was analyzed by RT-PCR and for both subtypes, the expression of all *cas* genes was greater during log growth than in stationary phase (Figure 4).

**Figure. 3:**
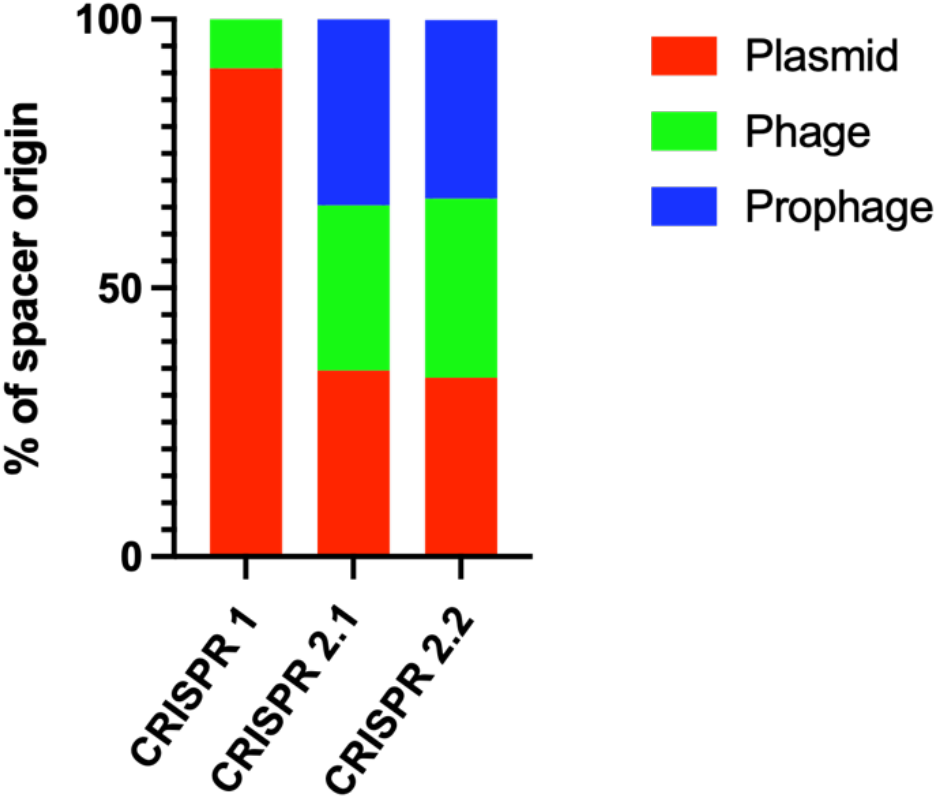
Origin of exogenous nucleic acid elements in the CRISP loci: Percentage of plasmids, phage and prophage DNA found in the spacer sequences for each of CRISPR array in Serratia Sp. Ag1.

**Figure 4.**
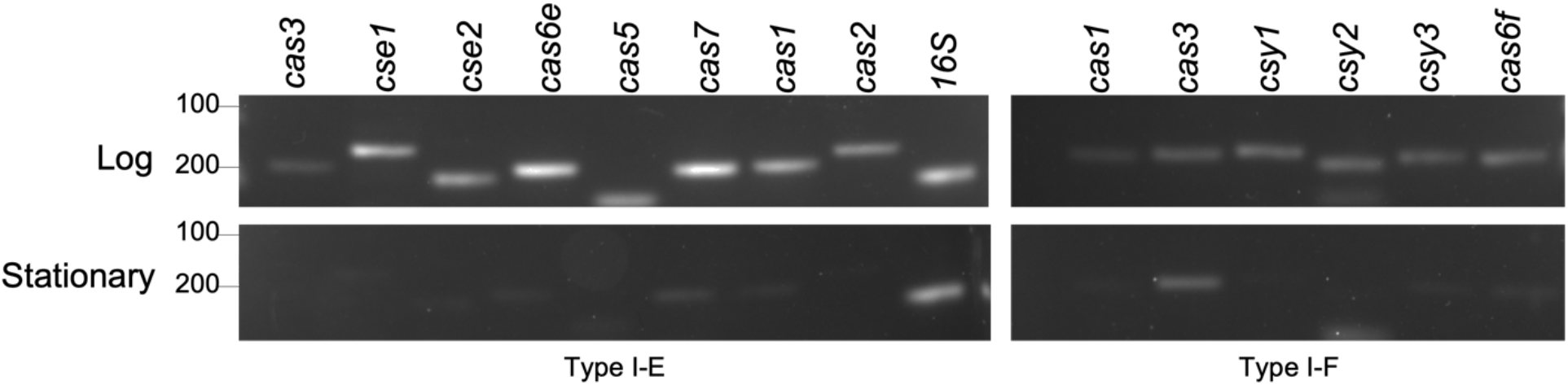
Expression of the Type I-E and Type I-F *cas* genes. The RT-PCR analysis of expression of cas and csy genes in logarithmic and stationary growth phase of Serratia sp Ag1.. A 100 base pair ladder was used, and sizes are indicated to the left of the gel images.

**Table 2:**
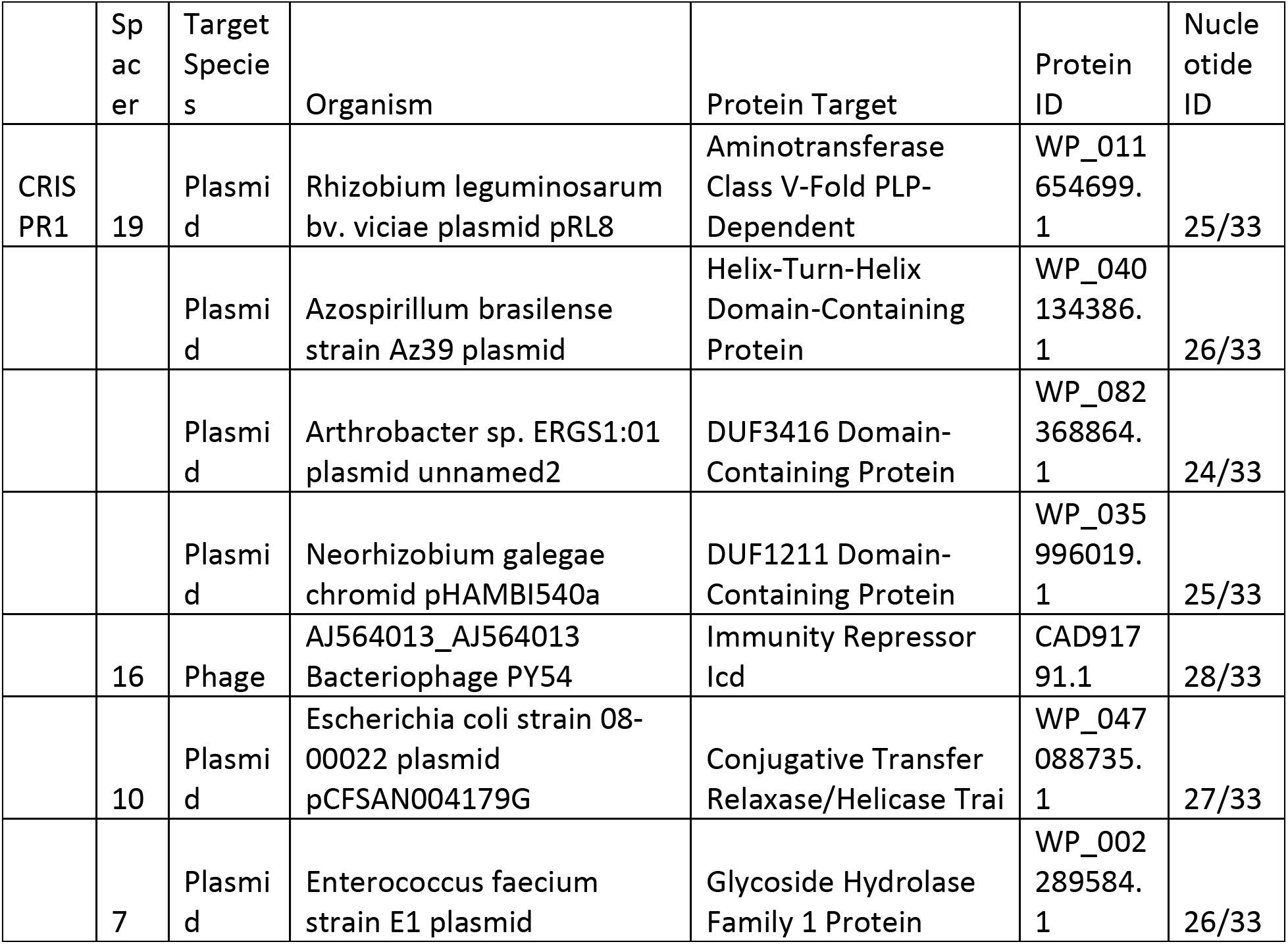
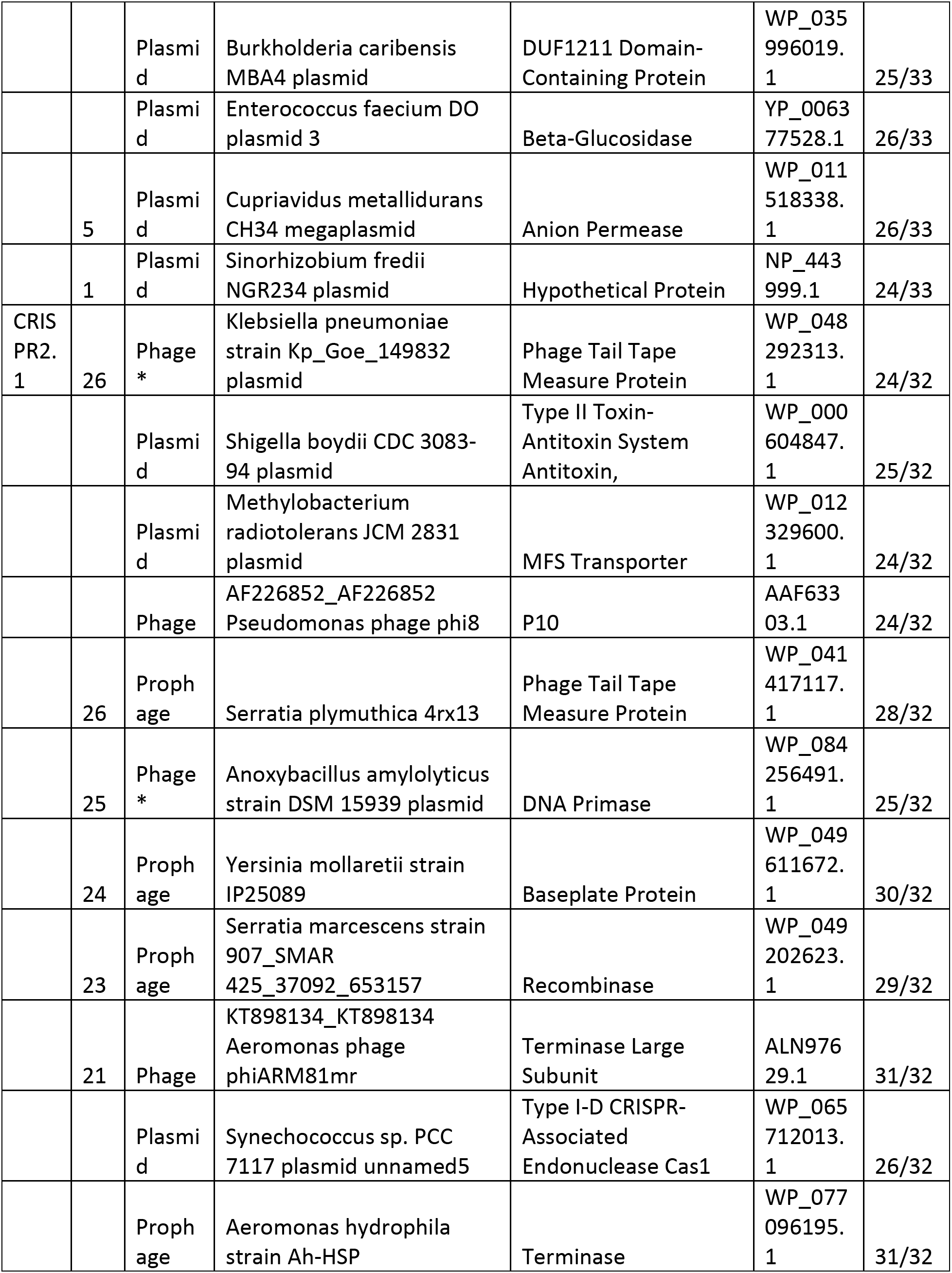
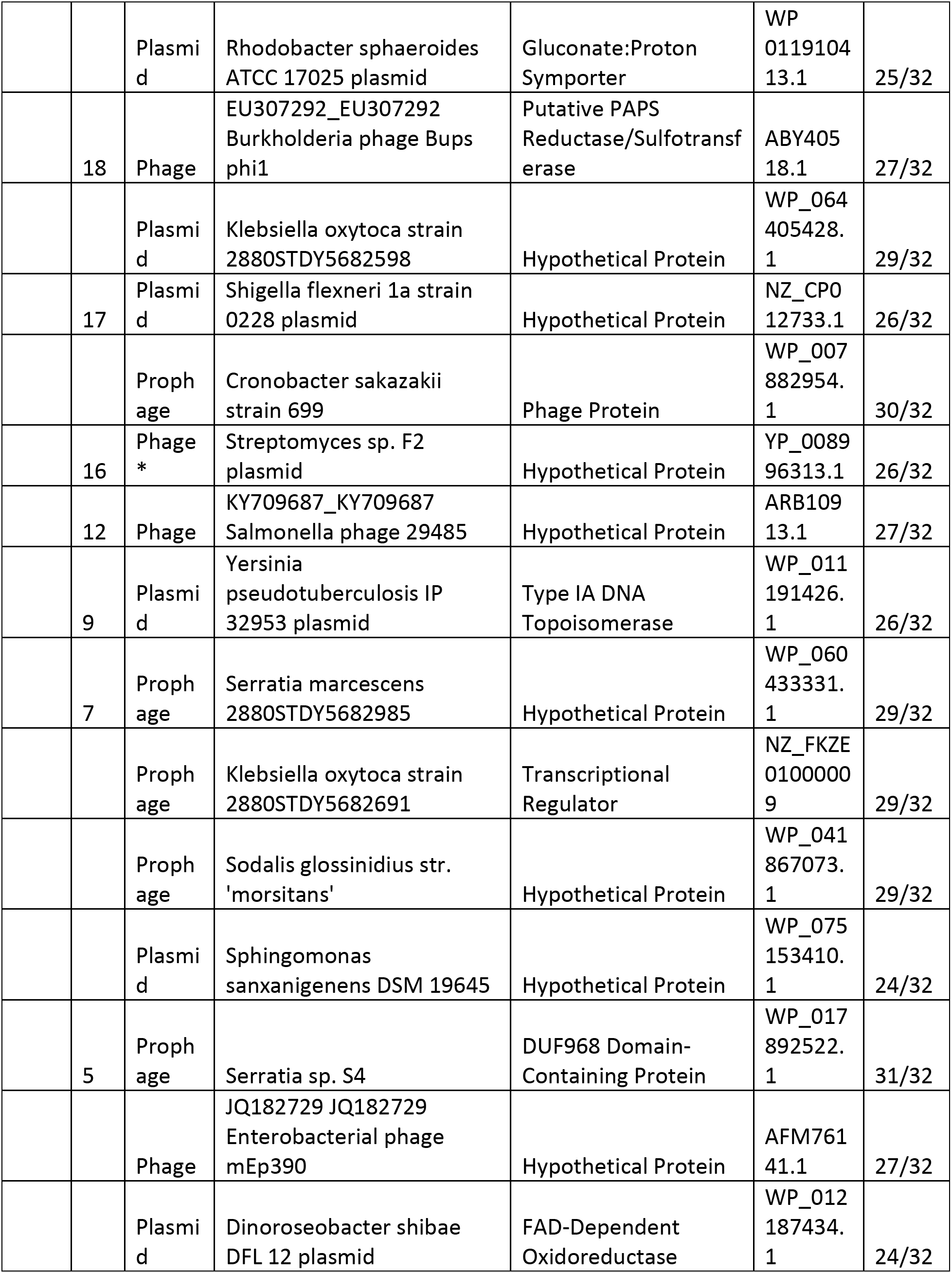
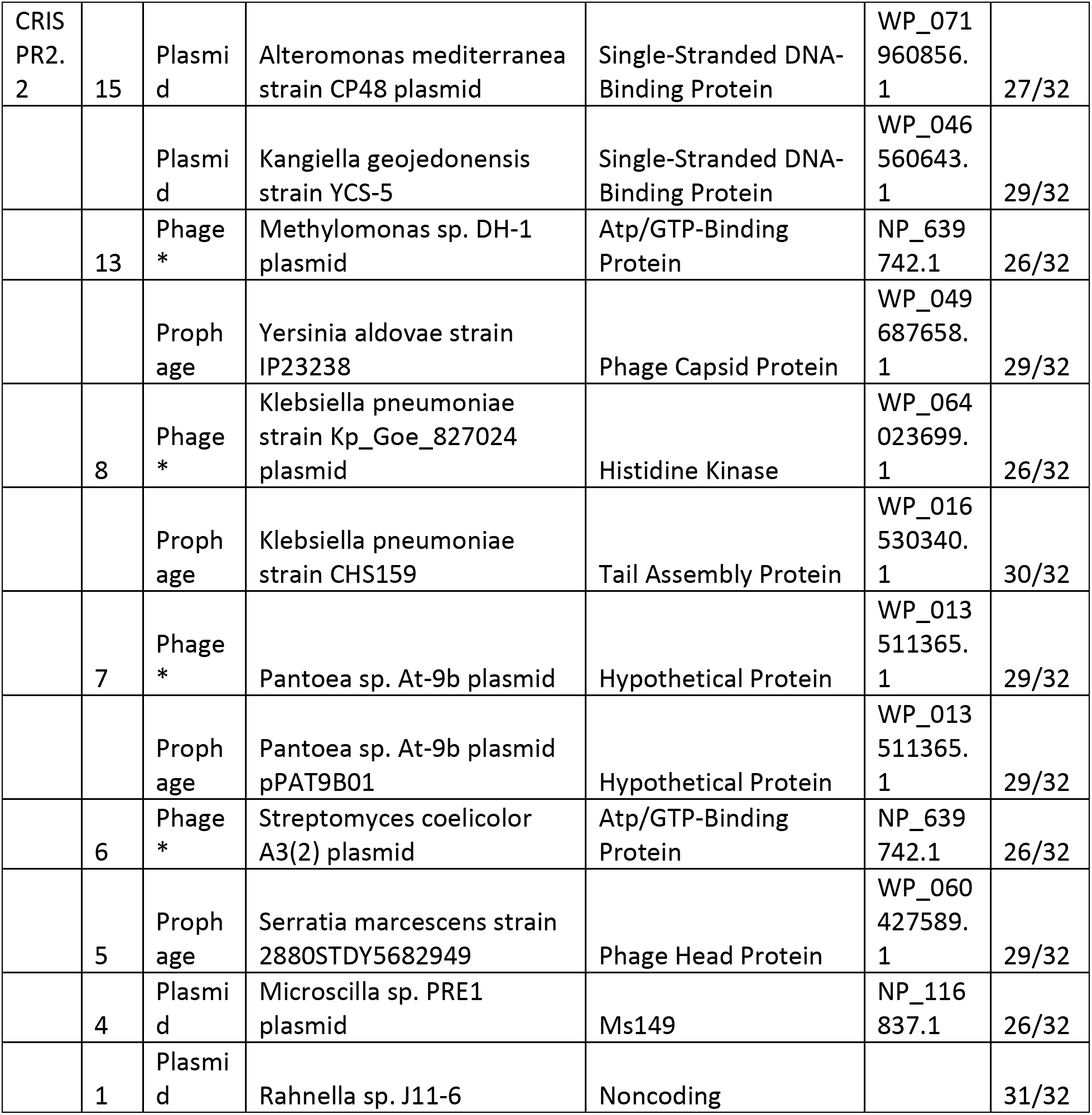
Protospacer identity: Origin of protospacer sequences matched against plasmids, phages and prophages.

## Discussion

Bacteria living in complex ecological settings are continuously challenged by predatory viruses. The CRISPR-cas adaptative immune systems of bacteria protect bacteria from some of these challenges by targeting foreign genetic material such as plasmids and bacteriophages. Here, we provide evidence of CRISPR cas system in the mosquito associated *Serratia* Sp. Ag1, which was isolated from *Anopheles gambiae* (39). We have identified two Type I CRISPR-cas systems, which are typically found in the *Enterobacteriaceae* family of bacteria (22, 27). CRISPR-Cas systems in *Serratia marcescens* have been previously described, and most strains harbor a Type I-F or both a Type I-E and Type I-F system (27, 34, 46, 47). One *Serratia* sp. (ATCC39006) contains both these Type I systems and also a Type III-A system (48).

Analysis of CRISPR spacer sequences in Ag1 confirmed the origin of many spacer sequences. Our results revealed 47% (23/49) of the spacer targets that we could identify originated from plasmids, while bacteriophage (phage and prophage) accounted for two thirds of the matched spacers. Overall, 53% (26/49) of spacers matched to phage or plasmid sequences. This is higher than other *Enterobacteriaceae* such as *Salmonella* (12%), *E. coli* (19%), 8% in STEC E. coli (Shuang) (49). That the newer spacers in the CRISPR2.1 and 2.2 arrays had matches to phage genomes that had been sequenced and deposited in NCBI suggests that these acquisition events have occurred relatively recently. Extensive spacer sequence analysis has been performed in the genomes of *Enterobacteriaceae* family bacteria that are human pathogens and commensals (27, 50–52). Discrete number of spaces from the *Enterobacteriaceae* members appear to be acquired from the extrachromosomal genetic elements such as plasmids and bacteriophages, while other spacers match to the bacterial host genome in non-prophage regions, however many remain of the spacer are of unknown origin.

Cas proteins play crucial roles in all three steps of CRISPR-Cas immunity (21, 53). Our results showed active expression of all *cas* genes during the actively dividing logarithmic growth phage of bacteria and attenuation of all but the Type I-F *cas3* gene during stationary phase. This is concordant with previous studies in *E. coli* showing repression of the Type I-E *cas3* gene expression during stationary phase compared to log phase (54–56). In the *E. coli* system, the transcription repressor, H-NS, is responsible for repressing *cas3* expression (54, 55).

CRISPR spacer sequences can be used for bacterial subtyping (57). The presence of the Type I-F cas3 in multiple *Serratia* spp. suggests that these genomes likely also contain CRISPR arrays. This would depend on CRISPR arrays being present in all strains of the species and to exhibit strain-to-strain variability that could be exploited for subtyping. Whole genome sequencing of four *Serratia marcescens* genomes showed that CRISPR-Cas systems were absent from half of these (34). The prevalence of CRISPR-Cas systems and the diversity of spacer content in other *Serratia* spp. is yet to be determined and would need to be performed to determine the utility of CRISPR typing in this bacterium.

While Cas1 and Cas2 are mainly involved in acquiring the spacers from newly invading phages and foreign genetic material, the cas2/3, csy complex are involved in the priming method of spacer acquisition (58, 59). Our results show the presence of newly acquired spacer sequences suggesting the adaptation is occurring actively in these bacteria. Hence, there is the possibility of recurrent encounter of phages with symbiotic bacteria in the mosquito gut. A recent study demonstrated that phage can alter bacterial levels in mosquitoes and alter their development in aquatic stages (60). These phages also may be part of the mosquito gut microbiome where they interaction with gut bacteria and compete for nutritional resources.

The CRISPR cas system in bacteria has been greatly explored in terms their application in different fields such as human and agriculture diseases (16, 21, 61, 62). However, analysis of CRISPR loci in the host associated symbiotic bacteria is limited, especially the role of CRISPR systems in the host-microbe interactions. Apart from anti-viral defence, CRISPR has been shown to be involved in the DNA repair, colonization, and host immune evasion (63–65). Hence, by modifying the CRISPR loci the colonization of bacteria in the host environment could be investigated. Such studies are important in deciphering the host-microbe interactions in complex ecological settings such as the mosquito microbiome. In this regard, further studies are needed to analyse CRISPR loci in the mosquito symbionts and understand mechanistic basis for CRISPR loci mediated host-microbiome interactions.

## Acknowledgments

We thank Naufa Amirani for help with the phylogenetic analyses of cas3 and Phillipe Horvath for providing the macro to visualize the spacer sequences.

## Author Contributions

H.E.R, S.H and N. S: conceptualization, methodology, investigation, data curation, writing – original draft preparation and visualization. H.E.R and N.S: investigation and data curation. H.E.R, S.H, G.L.H and N. S.: conceptualization methodology, data curation, resources, writing – review and editing and funding. G.L.H and N.S.: conceptualization, resources, writing – review and editing, supervision and funding.

## Reference

1. Hegde S, Rasgon JL, Hughes GL. The microbiome modulates arbovirus transmissionin mosquitoes. Current Opinion in Virology. 2015;15:97–102.

2. Cansado-Utrilla C, Zhao SY, McCall PJ, Coon KL, Hughes GL. The microbiome and mosquito vectorial capacity: rich potential for discovery and translation. Microbiome. 2021;9(1):111.

3. Gabrieli P, Caccia S, Varotto-Boccazzi I, Arnoldi I, Barbieri G, Comandatore F, et al. Mosquito Trilogy: Microbiota, Immunity and Pathogens, and Their Implications for the Control of Disease Transmission. Front Microbiol. 2021;12(633).

4. Huang W, Wang S, Jacobs-Lorena M. Use of Microbiota to Fight Mosquito-Borne Disease. Frontiers in Genetics. 2020;11.

5. Caragata EP, Tikhe CV, Dimopoulos G. Curious entanglements: interactions between mosquitoes, their microbiota, and arboviruses. Current Opinion in Virology. 2019;37:26–36.

6. Caragata EP, Short SM. Vector microbiota and immunity: modulating arthropod susceptibility to vertebrate pathogens. Curr Opin Insect Sci. 2022;50:100875.

7. Chandler JA, Liu RM, Bennett SN. RNA shotgun metagenomic sequencing of northern California (USA) mosquitoes uncovers viruses, bacteria, and fungi. Front Microbiol. 2015;6:185.

8. He X, Yin Q, Zhou L, Meng L, Hu W, Li F, et al. Metagenomic sequencing reveals viral abundance and diversity in mosquitoes from the Shaanxi-Gansu-Ningxia region, China. PLOS Neglected Tropical Diseases. 2021;15(4):e0009381.

9. Shi C, Beller L, Deboutte W, Yinda KC, Delang L, Vega-Rúa A, et al. Stable distinct core eukaryotic viromes in different mosquito species from Guadeloupe, using single mosquito viral metagenomics. Microbiome. 2019;7(1):121.

10. Atoni E, Wang Y, Karungu S, Waruhiu C, Zohaib A, Obanda V, et al. Metagenomic Virome Analysis of Culex Mosquitoes from Kenya and China. Viruses. 2018;10(1).

11. Shi C, Liu Y, Hu X, Xiong J, Zhang B, Yuan Z. A Metagenomic Survey of Viral Abundance and Diversity in Mosquitoes from Hubei Province. PLOS ONE. 2015;10(6):e0129845.

12. Batson J, Dudas G, Haas-Stapleton E, Kistler AL, Li LM, Logan P, et al. Single mosquito metatranscriptomics identifies vectors, emerging pathogens and reservoirs in one assay. eLife. 2021;10:e68353.

13. Kozlova EV, Hegde S, Roundy CM, Golovko G, Saldaña MA, Hart CE, et al. Microbial interactions in the mosquito gut determine Serratia colonization and blood-feeding propensity. The ISME Journal. 2020.

14. Hegde S, Khanipov K, Albayrak L, Golovko G, Pimenova M, Saldaña MA, et al. Microbiome interaction networks and community structure from laboratory-reared and field-collected *Aedes aegypti, Aedes albopictus*, and *Culex quinquefasciatus* mosquito vectors. Front Microbiol. 2018;9:715.

15. Grissa I, Vergnaud G, Pourcel C. The CRISPRdb database and tools to display CRISPRs and to generate dictionaries of spacers and repeats. BMC Bioinformatics. 2007;8:172.

16. Rath D, Amlinger L, Rath A, Lundgren M. The CRISPR-Cas immune system: Biology, mechanisms and applications. Biochimie. 2015;117:119–28.

17. Barrangou R. CRISPR-Cas systems and RNA-guided interference. Wiley Interdiscip Rev RNA. 2013;4(3):267–78.

18. Makarova KS, Koonin EV. Annotation and Classification of CRISPR-Cas Systems. Methods Mol Biol. 2015;1311:47–75.

19. Deveau H, Garneau JE, Moineau S. CRISPR/Cas System and Its Role in Phage-Bacteria Interactions. Annual Review of Microbiology. 2010;64(1):475–93.

20. Mohanraju P, Saha C, van Baarlen P, Louwen R, Staals RHJ, van der Oost J. Alternative functions of CRISPR-Cas systems in the evolutionary arms race. Nat Rev Microbiol. 2022;20(6):351–64.

21. Hille F, Charpentier E. CRISPR-Cas: biology, mechanisms and relevance. Philos Trans R Soc Lond B Biol Sci. 2016;371(1707):20150496.

22. Xue C, Sashital DG. Mechanisms of Type I-E and I-F CRISPR-Cas Systems in Enterobacteriaceae. EcoSal Plus. 2019;8(2).

23. Makarova KS, Zhang F, Koonin EV. SnapShot: Class 1 CRISPR-Cas Systems. Cell. 2017;168(5):946–.e1.

24. Makarova KS, Wolf YI, Alkhnbashi OS, Costa F, Shah SA, Saunders SJ, et al. An updated evolutionary classification of CRISPR-Cas systems. Nat Rev Microbiol. 2015;13(11):722–36.

25. Makarova KS, Haft DH, Barrangou R, Brouns SJ, Charpentier E, Horvath P, et al. Evolution and classification of the CRISPR-Cas systems. Nat Rev Microbiol. 2011;9(6):467–77.

26. Makarova KS, Wolf YI, Iranzo J, Shmakov SA, Alkhnbashi OS, Brouns SJJ, et al. Evolutionary classification of CRISPR-Cas systems: a burst of class 2 and derived variants. Nat Rev Microbiol. 2020;18(2):67–83.

27. Medina-Aparicio L, Dávila S, Rebollar-Flores JE, Calva E, Hernández-Lucas I. The CRISPR-Cas system in Enterobacteriaceae. Pathogens and Disease. 2018;76(1).

28. Cui L, Wang X, Huang D, Zhao Y, Feng J, Lu Q, et al. CRISPR-cas3 of Salmonella Upregulates Bacterial Biofilm Formation and Virulence to Host Cells by Targeting Quorum-Sensing Systems. Pathogens. 2020;9(1).

29. Munch PC, Franzosa EA, Stecher B, McHardy AC, Huttenhower C. Identification of Natural CRISPR Systems and Targets in the Human Microbiome. Cell Host Microbe. 2021;29(1):94–106 e4.

30. Hidalgo-Cantabrana C, Crawley AB, Sanchez B, Barrangou R. Characterization and Exploitation of CRISPR Loci in Bifidobacterium longum. Front Microbiol. 2017;8:1851.

31. Soto-Perez P, Bisanz JE, Berry JD, Lam KN, Bondy-Denomy J, Turnbaugh PJ. CRISPR-Cas System of a Prevalent Human Gut Bacterium Reveals Hyper-targeting against Phages in a Human Virome Catalog. Cell Host & Microbe. 2019;26(3):325–35.e5.

32. Crawley AB, Henriksen ED, Stout E, Brandt K, Barrangou R. Characterizing the activity of abundant, diverse and active CRISPR-Cas systems in lactobacilli. Sci Rep. 2018;8(1):11544.

33. Yang L, Li W, Ujiroghene OJ, Yang Y, Lu J, Zhang S, et al. Occurrence and Diversity of CRISPR Loci in Lactobacillus casei Group. Front Microbiol. 2020;11:624.

34. Scrascia M, D’Addabbo P, Roberto R, Porcelli F, Oliva M, Calia C, et al. Characterization of CRISPR-Cas Systems in Serratia marcescens Isolated from Rhynchophorus ferrugineus (Olivier, 1790) (Coleoptera: Curculionidae). Microorganisms. 2019;7(9).

35. Toyomane K, Yokota R, Watanabe K, Akutsu T, Asahi A, Kubota S. Evaluation of CRISPR Diversity in the Human Skin Microbiome for Personal Identification. mSystems. 2021;6(1).

36. Hegde S, Nilyanimit P, Kozlova E, Anderson ER, Narra HP, Sahni SK, et al. CRISPR/Cas9-mediated gene deletion of the ompA gene in symbiotic Cedecea neteri impairs biofilm formation and reduces gut colonization of Aedes aegypti mosquitoes. PLOS Neglected Tropical Diseases. 2019;13(12):e0007883.

37. Kistler KE, Vosshall LB, Matthews BJ. Genome Engineering with CRISPR-Cas9 in the Mosquito Aedes aegypti. CellReports. 2015;11(1):51–60.

38. Chaverra-Rodriguez D, Macias VM, Hughes GL, Pujhari S, Suzuki Y, Peterson DR, et al. Targeted delivery of CRISPR-Cas9 ribonucleoprotein into arthropod ovaries for heritable germline gene editing. Nature Communications. 2018;9(1):245.

39. Pei D, Hill-Clemons C, Carissimo G, Yu W, Vernick KD, Xu J. Draft Genome Sequences of Two Strains of Serratia spp. from the Midgut of the Malaria Mosquito Anopheles gambiae. Genome announcements. 2015;3(2):e00090–15-2.

40. Grissa I, Vergnaud G, Pourcel C. CRISPRFinder: a web tool to identify clustered regularly interspaced short palindromic repeats. Nucleic Acids Research. 2007;35(Web Server):W52–W7.

41. Horvath P, Romero Dennis A, Coûté-Monvoisin A-C, Richards M, Deveau H, Moineau S, et al. Diversity, Activity, and Evolution of CRISPR Loci in Streptococcus thermophilus. Journal of Bacteriology. 2008;190(4): 1401–12.

42. Biswas A, Gagnon JN, Brouns SJJ, Fineran PC, Brown CM. CRISPRTarget: bioinformatic prediction and analysis of crRNA targets. RNA Biol. 2013;10(5):817–27.

43. Kumar S, Stecher G, Tamura K. MEGA7: Molecular Evolutionary Genetics Analysis Version 7.0 for Bigger Datasets. Mol Biol Evol. 2016;33(7):1870–4.

44. Haft DH, Selengut J, Mongodin EF, Nelson KE. A guild of 45 CRISPR-associated (Cas) protein families and multiple CRISPR/Cas subtypes exist in prokaryotic genomes. PLoS Comput Biol. 2005;1(6):e60.

45. Pei D, Hill-Clemons C, Carissimo G, Yu W, Vernick Kenneth D, Xu J. Draft Genome Sequences of Two Strains of Serratia spp. from the Midgut of the Malaria Mosquito Anopheles gambiae. Genome Announcements.3(2):e00090–15.

46. Vicente CSL, Nascimento FX, Barbosa P, Ke H-M, Tsai IJ, Hirao T, et al. Evidence for an Opportunistic and Endophytic Lifestyle of the Bursaphelenchus xylophilus-Associated Bacteria Serratia marcescens PWN146 Isolated from Wilting Pinus pinaster. Microbial Ecology. 2016;72(3):669–81.

47. Srinivasan VB, Rajamohan G. Genome analysis of urease positive Serratia marcescens, co-producing SRT-2 and AAC(6’)-Ic with multidrug efflux pumps for antimicrobial resistance. Genomics. 2019;111(4):653–60.

48. Malone LM, Hampton HG, Morgan XC, Fineran Peter C. Type I CRISPR-Cas provides robust immunity but incomplete attenuation of phage-induced cellular stress. Nucleic Acids Research. 2021;50(1):160–74.

49. Dang TN, Zhang L, Zöllner S, Srinivasan U, Abbas K, Marrs CF, et al. Uropathogenic Escherichia coli are less likely than paired fecal E. coli to have CRISPR loci. Infect Genet Evol. 2013;19:212–8.

50. Hidalgo-Cantabrana C, Sanozky-Dawes R, Barrangou R. Insights into the Human Virome Using CRISPR Spacers from Microbiomes. Viruses. 2018;10(9):479.

51. Shariat N, Timme RE, Pettengill JB, Barrangou R, Dudley EG. Characterization and evolution of Salmonella CRISPR-Cas systems. Microbiology (Reading). 2015;161(Pt 2):374–86.

52. Yin S, Jensen MA, Bai J, Debroy C, Barrangou R, Dudley EG. The evolutionary divergence of Shiga toxin-producing Escherichia coli is reflected in clustered regularly interspaced short palindromic repeat (CRISPR) spacer composition. Appl Environ Microbiol. 2013;79(18):5710–20.

53. Sorek R, Lawrence CM, Wiedenheft B. CRISPR-Mediated Adaptive Immune Systems in Bacteria and Archaea. Annual Review of Biochemistry. 2013;82(1):237–66.

54. Westra ER, Pul Ü, Heidrich N, Jore MM, Lundgren M, Stratmann T, et al. H-NS-mediated repression of CRISPR-based immunity in Escherichia coli K12 can be relieved by the transcription activator LeuO. Mol Microbiol. 2010;77(6):1380–93.

55. Majsec K, Bolt EL, Ivančić-Baće I. Cas3 is a limiting factor for CRISPR-Cas immunity in Escherichia coli cells lacking H-NS. BMC Microbiology. 2016;16(1):28.

56. Mitić D, Radovčić M, Markulin D, Ivančić-Baće I. StpA represses CRISPR-Cas immunity in H-NS deficient Escherichia coli. Biochimie. 2020;174:136–43.

57. Shariat N, Dudley EG. CRISPRs: molecular signatures used for pathogen subtyping. Appl Environ Microbiol. 2014;80(2):430–9.

58. Swarts DC, Mosterd C, van Passel MWJ, Brouns SJJ. CRISPR Interference Directs Strand Specific Spacer Acquisition. Plos One. 2012;7(4).

59. Yosef I, Goren MG, Qimron U. Proteins and DNA elements essential for the CRISPR adaptation process in Escherichia coli. Nucleic Acids Res. 2012;40(12):5569–76.

60. Tikhe CV, Dimopoulos G. Phage Therapy for Mosquito Larval Control: a Proof-of-Principle Study. mBio. 2022;13(6):e0301722.

61. Liu Z, Dong H, Cui Y, Cong L, Zhang D. Application of different types of CRISPR/Cas-based systems in bacteria. Microbial Cell Factories. 2020;19(1):172.

62. Zhu H, Li C, Gao C. Applications of CRISPR–Cas in agriculture and plant biotechnology. Nature Reviews Molecular Cell Biology. 2020;21(11):661–77.

63. Varble A, Campisi E, Euler CW, Maguin P, Kozlova A, Fyodorova J, et al. Prophage integration into CRISPR loci enables evasion of antiviral immunity in Streptococcus pyogenes. Nature Microbiology. 2021;6(12):1516–25.

64. Babu M, Beloglazova N, Flick R, Graham C, Skarina T, Nocek B, et al. A dual function of the CRISPR-Cas system in bacterial antivirus immunity and DNA repair. Mol Microbiol. 2011;79(2):484–502.

65. Veesenmeyer JL, Andersen AW, Lu X, Hussa EA, Murfin KE, Chaston JM, et al. NilD CRISPR RNA contributes to Xenorhabdus nematophila colonization of symbiotic host nematodes. Mol Microbiol. 2014;93(5):1026–42.

